# Structural insights into the mechanism of phosphate recognition and transport by human XPR1

**DOI:** 10.1101/2024.08.19.608714

**Authors:** Wenhui Zhang, Yanke Chen, Zeyuan Guan, Yong Wang, Meng Tang, Zhangmeng Du, Jie Zhang, Meng Cheng, Jiaqi Zuo, Yan Liu, Qiang Wang, Yanjun Liu, Delin Zhang, Ping Yin, Ling Ma, Zhu Liu

**Affiliations:** National Key Laboratory of Crop Genetic Improvement, Hubei Hongshan Laboratory, Huazhong Agricultural University, Wuhan 430070, China; College of Life Sciences, Zhejiang University, Hangzhou, 310027, China

## Abstract

XPR1 is the only known protein responsible for transporting inorganic phosphate (Pi) out of cells, a function conserved from yeast to mammals. Human XPR1 variants lead to cerebral calcium-phosphate deposition, which are associated with a hereditary neurodegenerative disorder known as primary familial brain calcification (PFBC). Here, we present the cryo-EM structure of human XPR1 in both its Pi-unbound form and various Pi-bound states. XPR1 features 10 transmembrane α-helices that form an ion channel-like architecture. Multiple Pi recognition sites are arranged along the channel, facilitating Pi ion transport. Two arginine residues, subject to pathogenic mutation in PFBC families, line the translocation channel and serve to bind Pi ion. Clinically linked mutations in these arginines impair XPR1’s Pi transport activity. To gain dynamic insights into the channel-like transport mechanism, we conducted molecular dynamics simulations. The simulations reveal that Pi ion undergoes a stepwise transition through the sequential recognition sites during the transport process. Together with functional analyses, our results suggest that the sequential arrangement of Pi recognition sites likely enable XPR1 to use a “relay” process to facilitate Pi ion passage through the channel, and they establish a framework for the interpretation of disease-related mutations and for the development of future therapeutics.

**One Sentence Summary:** Combined cryo-EM, molecular dynamics simulations and functional studies demonstrate that human XPR1 employs a channel-like transport mechanism to export inorganic phosphate out of cells

## Introduction

Inorganic phosphate (Pi) is essential for life, used wildly in biomolecules synthesis, cell metabolism and energy supply. To nourish cell, humans uptake Pi through Na^+^-coupled Pi importers, which belong to the solute carrier 20 (SLC20) and SLC34 transporter families^1–5^. In parallel, cell exports intracellular Pi to regulate Pi homeostasis and prevent the cytotoxicity caused by Pi accumulation. Elevated Pi levels cause major disorders, with severe biochemical and clinical consequences^6–8^.

XPR1 is the only known exporter that transports Pi out of cell in humans^9^, expressed in all tissues^10^. It is a passive transporter belonging to the SLC53 family. Unlike active Pi importers, which utilize energy from the movements of co-translocated ions (eg., Na^+^ or H^+^) to transport Pi into cells^11–13^, XPR1 facilitates Pi efflux independently of co-translocated ions^14–16^. Loss-of-function variants of XPR1 accumulate cytosolic Pi and lead to cerebral calcium-phosphate deposition, which are associated with a genetic disease known as primary familial brain calcification (PFBC)^17–20^. PFBC is an adult-onset neurodegenerative disorder characterized by a wide range of clinically heterogeneous symptoms, including dystonia, parkinsonism, dementia, depression, chorea and others^21^. Currently, there are no targeted drugs or specific treatments available^22,23^. XPR1 is abundant in human platelets, where inhibiting its transport activity increases the risk of thrombosis in mouse models^24^. XPR1 deficiency in ovarian cancer cell causes toxic accumulation of cytosolic Pi, leading to cell death^25^. Dysregulation of Pi efflux could thus present a new, potential strategy for anticancer therapy^16,25^. Given the significance and pathomechanisms of XPR1, it presents a promising therapeutic target for diseases and cancers^17,25–27^. However, the molecular basis for this fundamental transporter remains unknown. We therefore set out to determine the cryo-electron microscopy (cryo-EM) structure of human XPR1, and to establish how it recognizes and transports Pi across cell membrane.

## Results

### The overall cryo-EM structure

XPR1 contains a transmembrane domain and a cytoplasmic N-terminal SPX (named after yeast SYG1, Pho81 and human XPR1) domain. The transmembrane domain is responsible for facilitating Pi efflux, and the SPX domain is thought to modulate this function by binding inositol pyrophosphate (PP-InsP) stimuli^9,14,28–30^. To aid cryo-EM structure determination, we prepared XPR1 with the presence of InsP_6_ (Fig. S1, and Methods). InsP_6_ is a commercially available surrogate for PP-InsPs^31,32^, which has been indicated to potentially limit the structural heterogeneity of SPX domains^33,34^. A total of 20,420 cryo-EM micrographs were collected and processed through single-particle analysis (Fig. S2, and Methods). Following series 3D classification, 1,168,388 particles were pooled, capturing millions of snapshots of the molecules in their various conformational states. We then reprocessed this particle pool into 10 classes using heterogeneous cryo-EM structure reconstruction, ensuring that each particle was assigned to only one class. Consequently, four distinct density maps of XPR1 were obtained from the final 10 reconstructions, with overall resolutions ranging from 2.9 to 3.3 Å. The highest resolution map is identified as the Pi-unbound form, while the other three maps correspond to Pi-bound forms, each displaying additional nonprotein densities within a membrane-spanning pore of the XPR1 transmembrane domain (Discussed later). The EM density of SPX domain was observed to be poor (Fig. S3A), probably owing to its mobility. By contrast, the density map of the transmembrane domain was well resolved (Fig. S3A), enabling the construction of an atomic model (Fig. S3B, C, and Table S1). XPR1 forms dimers in the micelles (Fig. S3A, B), with transmembrane domains loosely contacted, indicating that they may function as independent units.

The transmembrane domain of XPR1 contains 10 transmembrane α-helices (TM1-10) and folds into two structurally distinct sub-domains, with N and C termini on the intracellular side (Fig. 1). We refer to the N-terminal portion as N domain, that is formed by TM1-TM5 and a short amphipathic helix (AH) lying parallel to the membrane. The C-terminal portion harbors the conserved EXS (named for homologous regions found in yeast ERD1 and SYG1 and human XPR1) domain^35,36^ (Fig. S4), that is made up of TM6-TM10. The EXS domain associates with TM5, creating a pore that spans across the membrane (Fig. 1A). Notably, the TM9 bends and positions TM9b close to the central pore axis on the extracellular side (Fig. 1A), suggesting its potential role in Pi export. This experimental structure differs from the predicted model of AlphaFold2^37^ (Fig. S5), and a 3-dimensional structural homology search with the program DALI^38^ found no similar known structures, indicating a specific mechanism for phosphate recognition and transport in XPR1.

**Fig. 1.**
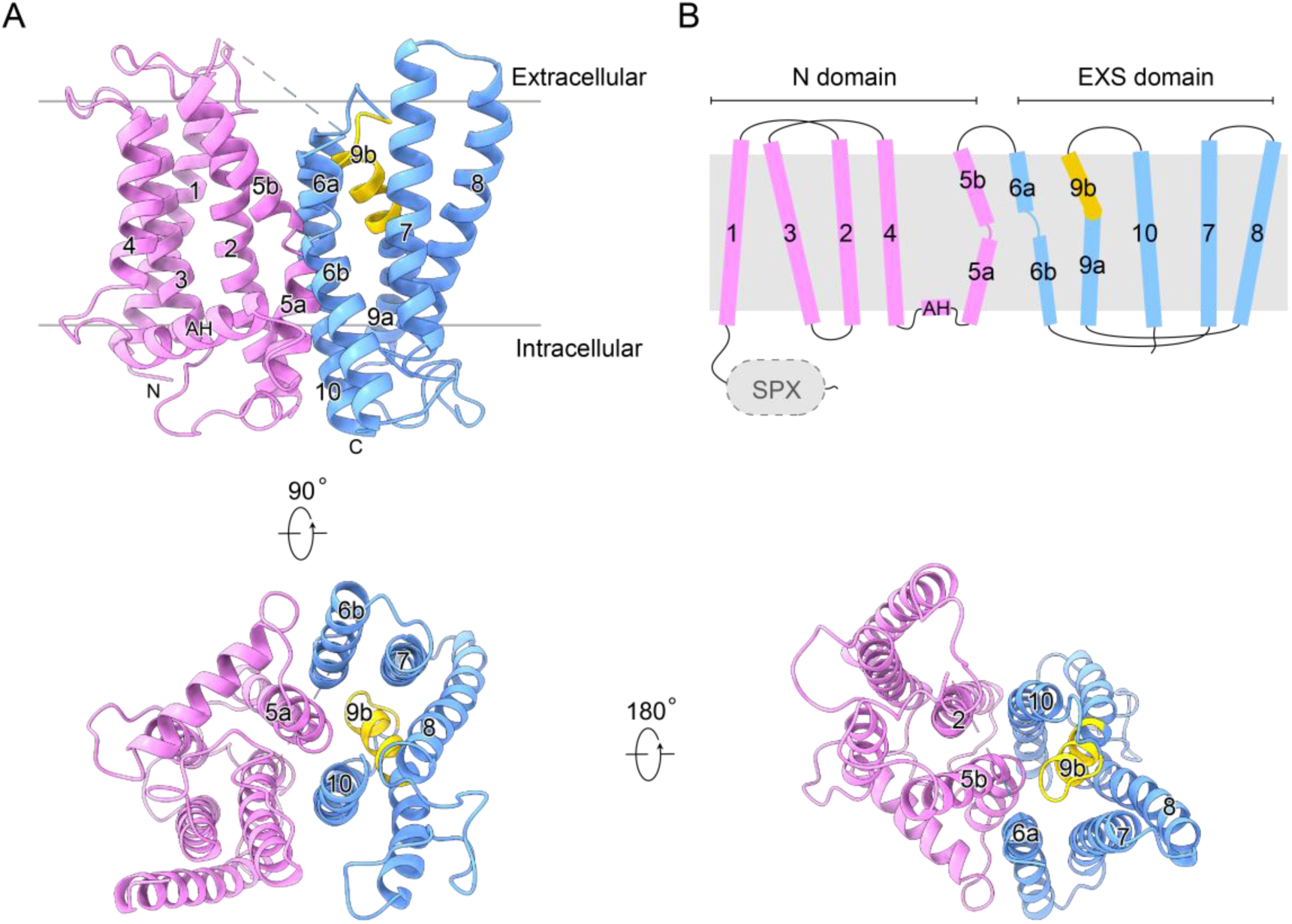
Structure of XPR1 transmembrane domain. **A** Cartoon representation of the structure. The N domain and EXS domain are colored in magenta and blue, respectively. TM9b is colored in yellow. **B** Schematic topology diagram of the structure. The gray background indicates the membrane bilayer. The N-terminal SPX domain of XPR1 is invisible in the determined cryo-EM structure.

### The structure basis for phosphate recognition and transport

To gain insight into the pathway for Pi ion translocation, we conducted CAVER^39^ analysis for the resolved structure. The analysis revealed a continuous, solvent-accessible pathway within the XPR1 transmembrane domain, featuring an ion channel-like architecture (Fig. 2A, B, and Fig. S6A). The narrowest point of this channel has a radius of approximately 1.2 Å (Fig. 2B), which exceeds the water access limit (1 Å)^40^, suggesting its permeability to solvents and potential for Pi ion conduction. As this channel is smaller than the ionic radius of Pi (2.38 Å), the observed conformation is presumably a partially open state. The intracellular entrance of the translocation channel is formed by TM10, TM6b, TM7, TM5a and TM8 (Fig. 2A, Fig. S6A). Owing to the bending of TM9, the channel is curved and created by TM9b, TM10, TM6a, TM5b and the tip of TM2 on the extracellular side. The channel is primarily lined with polar residues, characterized by a positively charged interior wall (Fig. 2A, Fig. S6B), which presumably facilitates the permeation of anionic Pi ions. The narrow regions along the ion permeation pathway is constricted mainly by residues D398, Y483, N401, R570, S404 and W573 (Fig. 2B), implying their roles in Pi transport.

**Fig. 2.**
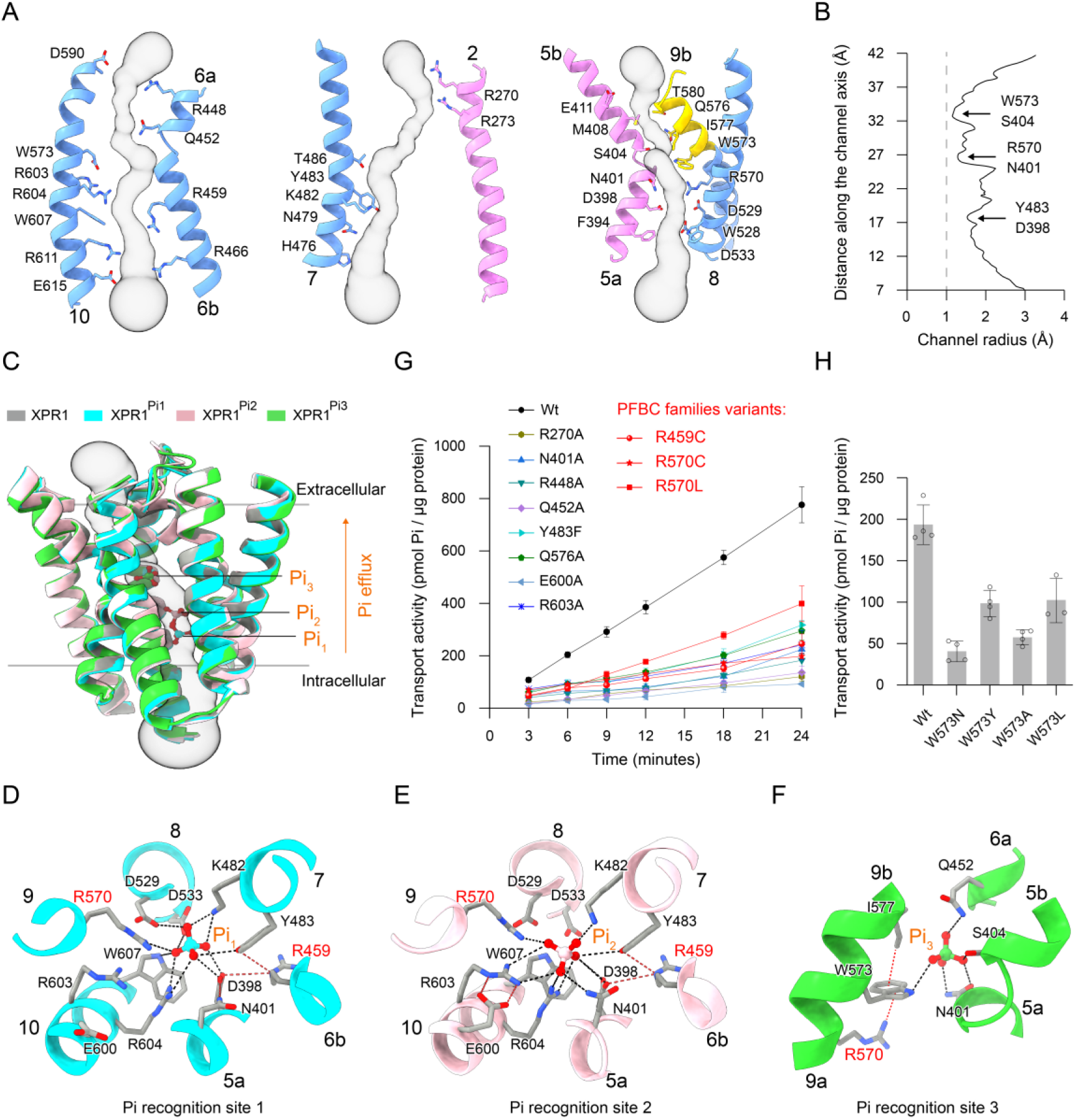
Phosphate recognition and transport by XPR1. **A** Channel-like architecture. Gray surface represents the channel path within XPR1 transmembrane domain. Transmembrane α-helices forming the channel are indicated, and channel-lining residues are shown as sticks. **B** Pore radius along the channel. The gray dashed line indicates a radius of 1 Å for water access limit. **C** Structures superposition. The transmembrane domain structures of Pi-unbound XPR1, Pi-bound XPR1^Pi1^, Pi-bound XPR1^Pi2^ and Pi-bound XPR1^Pi3^ are colored in gray, cyan, pink and green, respectively. Only the channel-forming transmembrane α-helices are represented for clarity. Pi ion in each structure is shown as sticks. **D-F** Pi recognition sties 1, 2 and 3, observed in the structures of XPR1^Pi1^, XPR1^Pi2^, XPR1^Pi3^, respectively. The black dashed lines indicate plausible interactions between the Pi ion and the protein, while the red dashed lines represent the interactions that may indirectly contribute to Pi coordination. **G-H** Activity assay. The Pi transport activity is determined by measuring phosphate uptake into liposomes containing wild-type and variant forms of XPR1. Activity of PFBC families variants, R459C, R570C and R570L, are highlighted in red. The time-course activity (**G**) was collected at external Pi concentration of 500 μM, with transport reaction times of 3, 6, 9, 12, 18 and 24 minutes. The single-point activity (**H**) was collected at the external Pi concentration of 500 μM, with a transport reaction time of 6 minutes. The data presented are the average of 3-4 independent assays, each with technical triplicates. The error bars indicates the SD.

In parallel to the above-resolved XPR1 transmembrane domain structure (Pi-unbound XPR1), our structural heterogeneous analysis has reconstructed three additional cryo-EM maps with resolutions of 3.1-3.3 Å (Fig. S2, and Methods). These reconstructed maps revealed extra nonprotein densities positioned at various locations inside the translocation channel. These clear densities were putatively modeled with Pi ions and are well-positioned by surrounding residues within the channel (Fig. 2C-F, Fig. S7). As no phosphate was added during our sample preparation, we assume that the observed Pi ions were copurified with XPR1 endogenously. Moving in the intracellular-to-extracellular direction, the three structures are denoted as XPR1^Pi1^, XPR1^Pi2^ and XPR1^Pi3^ based on the positions of Pi ions lined along the channel (Fig. 2C). The three structures are also dimeric and their protomers show overall structural similarity to that of Pi-unbound XPR1, with RMSD of 0.26-0.62 Å. This suggests that Pi binding does not induce large conformational changes. The aqueous pore within XPR1 transmembrane domain (Fig. 2B), together with aligned Pi ions along the permeation pathway (Fig. 2C), suggests that XPR1 may facilitate the passive diffusion of Pi ion through this channel without requiring large conformational changes, as observed in Pi importers. Pi importers allow only one side to be solvent-associable at a time. They exhibit an outward-open conformation for Pi uptake from the extracellular side and subsequently undergo large conformational changes to adopt an inward-open conformation, allowing for Pi release into cytosol^11–13^. This conformational transition is driven by the energy derived from movements of other co-translocated ions (e.g., Na^+^ or protons) down their concentration gradients^11–13^. By contrast, the Pi export capability of XPR1 is independent of pH gradient across the cell membrane^14,15^, and no co-translocated ions have been found^16^. These previously observed phenomena are now supported and aligned with the channel-like architecture of XPR1.

Our structures reveal stepwise states of Pi ion transitions within the translocation channel, providing a framework for understanding how XPR1 recognizes and transports phosphate. The translocation channel is lined by a set of polar and charged residues, which provide sites for the recognition of ionic Pi (Fig. 2A, Fig. S6B). At Pi recognition site 1, the structure of XPR1^Pi^^1^ reveals that the Pi ion (Pi_1_) is primarily recognized through interactions with the side chains of D398, K482, Y483, D529, D533, R570, R604, and the indole N atom of the W607 (Fig. 2D). These interactions include hydrogen bonds and salt bridges between the Pi_1_ ion and the protein, with details listed in Table S2. Additionally, R459 and N401 on the periphery likely orient the side chains of D398 and Y483, facilitating their coordination of Pi_1_ at the binding site. Subsequently, the Pi ion further moves from the recognition site 1 to site 2, disengaging from D529, D533 and W607, and forming new interactions with R603 and N401 (Fig. 2E, Table S2). During this transition, the peripheral E600 rotates and establishes salt bridges with R603, positioning R603 for Pi coordination. When the Pi ion moves upward (intracellular-to-extracellular direction), the XPR1^Pi3^ structure captures a state where the Pi ion is coordinated at recognition site 3 (Fig. 2F). At this site, the Pi ion dissociates from R570 and forms direct interactions with the side chains of N401, S404, Q452 and W573. Additionally, R570 and I577 contribute indirectly to this recognition site by forming cation-π and CH-π interactions with W573, respectively. These observed Pi-recognizing residues are largely conserved among EXS domain-containing proteins across different species (Fig. S4), suggesting functional and mechanistic conservations.

To gain functional support for the above-identified key residues in Pi transport, we reconstituted XPR1 into liposomes and performed Pi transport assays (Methods). Results showed that the reconstituted XPR1 exhibited Pi transport activity and transported Pi into the proteoliposomes in an external Pi concentration-dependent manner (Fig. S8), as well as a time-dependent manner (Fig. 2G). The Pi transport activity correlates linearly to the external Pi concentration up to 1 mM in the established assays (Fig. S8), implying that XPR1 may operate as a low-affinity Pi transporter with a *K*_m_ in millimolar range. By substituting the Pi-recognizing residues D398, K482, D533, R604, R603, N401 and Q452 with alanine, and Y483 and W607 with phenylalanine, we could purify the Y483F, R603A, N401A and Q452A in sufficient quantities to perform the proteoliposome-based transport assays. Time-course results showed that XPR1 carrying Y483F, R603A, N401A or Q452A substitution has impaired Pi transport activity (Fig. 2G), revealing crucial roles of these residues in Pi transport. Moreover, consistent with the structural role of the E600 side chain in positioning R603 at Pi recognition site 2 (Fig. 2E), replacing E600 with alanine resulted in a reduced Pi transport kinetic of XPR1 (Fig. 2G). At Pi recognition site 3, the polar NE1 atom of W573 and its bulky indole ring likely cooperate in Pi coordination (Fig. 2F). Supporting this, substituting the side chain of W573 with polar or hydrophobic groups such as W573N, W573Y, W573A and W573L reduced the Pi transport activity of XPR1 (Fig. 2H).

Additionally, we evaluated the significance of these Pi-recognizing residues by performing Pi transport assays in a heterologous yeast system. We utilized a yeast mutant strain (YP100) lacking Pi transporters, which cannot grow on YNB media^41^. Previous study found that transformation of this mutant yeast with rice OsPHO1;2, a functional homologue of human XPR1, can rescue the yeast growth phenotype^15^. Similarly, we observed that complementation with XPR1 also restored yeast growth (Fig. S9). In contrast, yeast strains complemented with XPR1 mutants, including N401A, Q452A, Y483F, W573A, W573N, W573L, W573Y, E600A and R603A, exhibited impaired growth (Fig. S9). These findings corroborate our proteoliposome-based assays results (Fig. 2G, H), underscoring the crucial roles of these residues in Pi transport.

### Clinically linked residues in XPR1 function in phosphate recognition and transport

*XPR1* responsible for autosomal dominant inheritance is a causative gene identified in primary familial brain calcification (PFBC) families^17,19,20^, and the hereditary R459C, R570C and R570L missense variants have been found to be pathogenic^18,23,42–44^. Our structures reveal that R459 contributes to establish the Pi binding network at recognition sites 1 and 2, while R570 plays a direct role in Pi recognition during its translocation (Fig. 2D-F). Substituting the side chains of R459 and R570 might perturb Pi binding and potentially impair XPR1 activity. Indeed, we find that the PFBC families variants, R459C, R570C and R570L, exhibit a significant reduction in Pi transport activity (Fig. 2G). The impaired function of these mutants is also evident in heterologous yeast assays (Fig. S9). This Pi transport deficiency observed in these hereditary XPR1 variants aligns with clinical findings of accumulated cytosolic Pi and cerebral calcium-phosphate deposition in PFBC patients. Therefore, our structure and function analyses provide mechanistic insights into the correlation between patient mutations and XPR1 function.

### Dynamic insights into phosphate transport mechanism

To gain dynamic insights into the channel-like transport mechanism of XPR1, we performed two independent 1000-ns all-atom molecular dynamics (MD) simulations using the Pi-unbound XPR1 cryo-EM structure (Methods). These simulations aimed to assess the stability of the structure and evaluate the permeation properties of the channel-like pore. The simulations results showed that the overall structure remained stable, as reflected by the relatively small root mean square deviations (RMSD) of the transmembrane helices, despite fluctuations in the loop regions (Fig. S10A-C). Throughout the simulations, the transmembrane pore consistently exhibited hydration (Fig. S10D), suggesting its potential to facilitate Pi ion permeation.

To further investigate the Pi transport process, we performed MD simulations on the Pi-bound XPR1 (XPR1^Pi1^) structure under an applied electric field (Methods). Following the simulations, we observed an export event of Pi ion through the channel, where the Pi ion dissociated from the recognition site 1, moved upward, and released into the extracellular solution (Supplementary Video 1). Minor structural changes were observed in XPR1 during the transport process, further supporting its channel-like properties (Fig. S11A). The simulations also showed that the transmembrane pore remained hydrated, similar to the Pi-unbound form, allowing the Pi ion to remain solvated and diffuse freely within the channel (Fig. 3A). Interestingly, the Pi ion traversed the entire length of the channel within a relatively short time, following a stepwise mechanism that involved three distinct binding sites (Fig. 3B, Fig. S11B, and Supplementary Video 1). These binding sites aligned well with the conformations observed in our cryo-EM structures, suggesting they likely represent stable intermediates in the transport process.

**Fig. 3.**
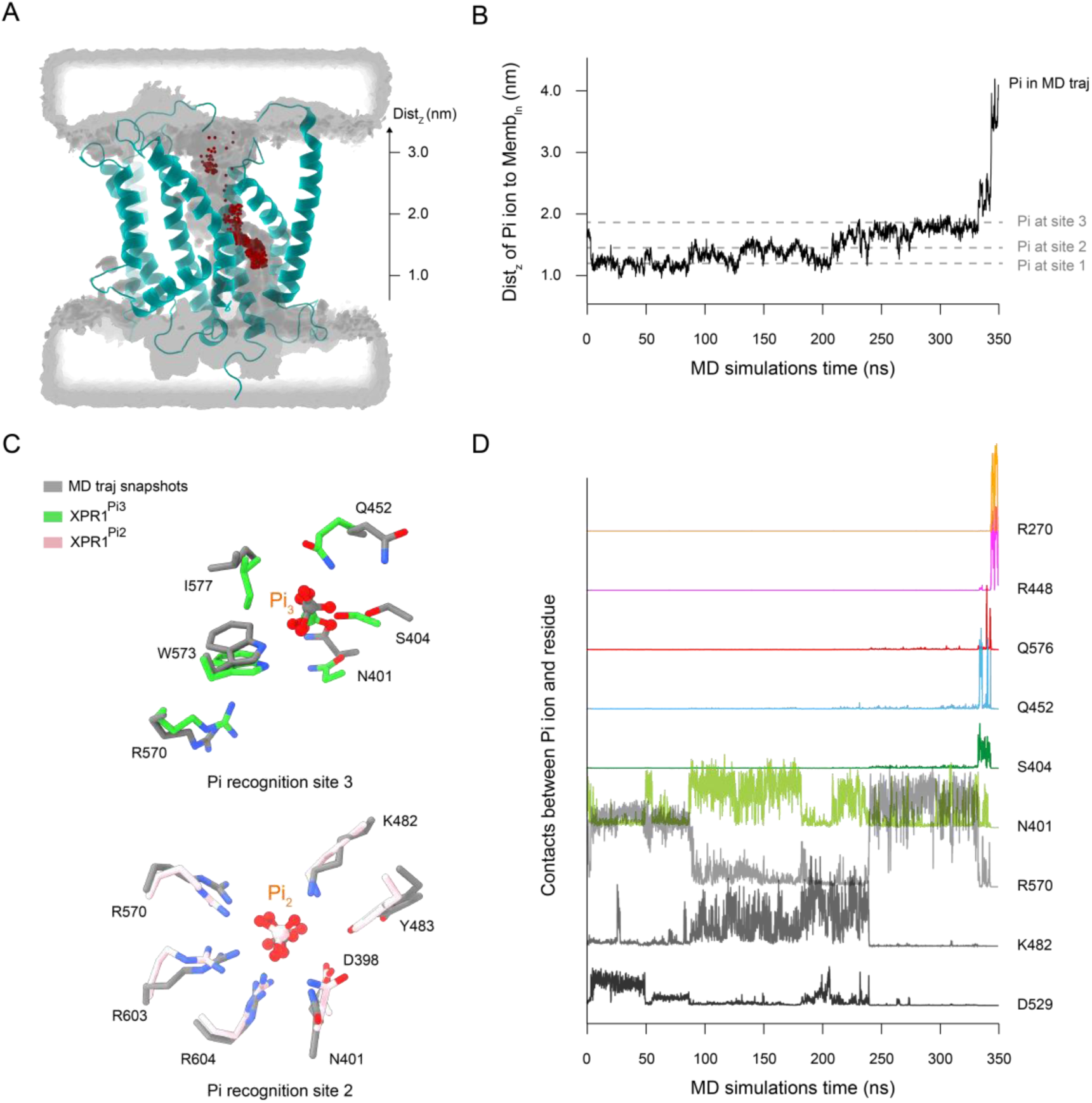
Molecular dynamics simulations of Pi ion translocation through XPR1. **A** Side-cut view of the channel, showing the average water density from the simulation as a gray transparent surface. The Pi ion positions sampled across all simulation frames are depicted as red spheres, illustrating its diffusion within the translocation channel. The distance (Dist_z_) between the Pi ion and the center of mass of the phosphate head groups of lipids in the inner leaflet along the membrane’s normal direction (Z-axis) is defined and labeled on the right. **B** Movement of the Pi ion through the channel, monitored by Dist_z_ in the simulations. The Pi ion is released into the extracellular solution at approximately 350 ns of simulation time. The gray dashed lines represent the positions of Pi ions as observed in the cryo-EM structures of XPR1^Pi1^, XPR1^Pi2^, and XPR1^Pi3^. **C** Comparison between cryo-EM structures and conformational snapshots from MD simulations. The Pi recognition sites 2 and 3 conformations in the XPR1^Pi2^ and XPR1^Pi3^ structures are colored pink and green, respectively, while similar conformations captured at ∼138 ns and ∼333 ns of simulation time are shown in gray. **D** Trajectories of Pi-residue contacts to show representative interactions between the Pi ion and specific residues during its transport process.

Our cryo-EM structures provide insights into the mechanism by which the Pi ion transitions from the site 1 to site 3 (Fig. 2D-F). Specifically, as the Pi ion moves upward, it first dissociates from residues D529, D533 and W607 at site 1 (Fig. S11C), then forms a new interaction with R603 at site 2 before transitioning to the site 3, where it is further recognized by residues S404, Q452 and W573 (Fig. S11C). Supporting this mechanistic pathway, MD simulations revealed that the Pi ion undergoes a stepwise transition through D529, R603, and Q452, corresponding to recognition sites 1, 2 and 3, respectively (Fig. S11D). Notably, despite beginning with the XPR1^Pi1^ structure, the simulations sampled conformations similar to those experimentally resolved structures of XPR1^Pi2^ and XPR1^Pi3^ (Fig. 3C). Throughout the simulations, the movement of Pi ion was facilitated by orchestrated interactions with residues from recognition site 1 to site 3 (Fig. 3D). Moreover, as the Pi ion disengaged from residues in site 3 (N401, S404, and Q452), it interacted with Q576, R448 and R270 (Fig. 3D) which are positioned above the site 3 and near the channel exit (Fig. 2A). These findings suggest that these residues may facilitate the Pi ion’s exit from the channel. Consistent with this, XPR1 mutants Q576A, R448A and R270A exhibit reduced Pi transport activity (Fig. 2G, Fig. S9). Together, these observations reinforce our structural findings of XPR1 and enhance the understanding of the channel-like transport mechanism. We propose that XPR1 likely employs a “relay” process to facilitate Pi ion passage through the channel, with sequential Pi recognition sites arranged along its pathway.

## Discussion

Elevated levels of cytosolic phosphate are cytotoxic due to its presence as a potent metal chelator and a pervasive inhibitor of cellular enzymes. XPR1 and its evolutionarily conserved orthologues are the proteins that transport cytosolic phosphate out of cells^9,15,45–47^. A very recent study discovered that the XPR1 orthologue in fruit fly (Pxo) lowers cellular phosphate levels, by generating a newly defined phosphate-storing organelle (PXo body) and transporting cytosolic phosphate into the PXo body^48^. These transporters belong to the structurally uncharacterized SLC53 family. Here the experimental structure of XPR1 transmembrane domain, together with its multiple functional states bound to phosphate, provides the first snapshots for understanding the structure-function relationships of SLC53 family transporters.

XPR1 mutations are linked to PFBC neurodegenerative disorder for which there are currently no targeting drugs and specific treatments available^17,21–23^. Lack of structural and related functional understanding limits patient mutation interpretation to disease mechanism. In this study, our structure and function analyses reveal that the pathogenic mutation residues (R459 and R570) line the phosphate translocation channel and act in phosphate recognition and transport. Notably, the MD simulations indicate that residue R570 interacts with Pi ion throughout the entire transport process (Fig. 3D), underscoring the significance of R570. These mechanistic understanding might provide an opportunity for targeting XPR1 in the development of future therapeutics.

XPR1 exhibits a channel-like architecture that facilitates phosphate efflux, distinguishing it from phosphate importers that use an alternating access mechanism for phosphate uptake^11,12,49,50^. This variety in mechanisms for balancing cellular phosphate levels offers cell regulatory layers. The activity of phosphate importers is regulated by the concentration gradient of other co-translocated ions (e.g., Na^+^ or proton)^11–13,16^, but no co-translocated ions are required for XPR1 function^14–16^. Instead, XPR1 harbors an additional cytoplasmic SPX domain, which modulates XPR1 function by sensing PP-InsP nutrient messengers^9,14,28–30^. PP-InsPs have emerged as cellular high-Pi signals in regulating phosphate homeostasis^31–33,51,52^. The limited cryo-EM density observed for the SPX domain in our reconstructed map suggests its mobility. This mobile nature might facilitate PP-InsPs targeting and regulation. Further dynamic studies are needed to understand the regulatory role and mechanism of XPR1 SPX domain.

## Methods

### Protein expression and purification

Human XPR1 DNA was subcloned into a pMlink vector encompassing a C-terminal 3×Flag tandem affinity tag. Point mutations were introduced into XPR1 genes by overlapping PCR and were verified by DNA sequencing. Proteins were expressed in Expi293F^TM^ cells (Invitrogen) by transient transfection. Cells grown in Union-293 media (Union-Biotech, Shanghai) were transfected with linear polyethyleneimine (PEI) (Polysciences) at a cell density of 2.0×10^6^ cells per ml^-1^. The transfected cells were cultured for another 60 hours before harvesting.

To prepare the cryo-EM sample, cultured cells were collected and resuspended in the TBS buffer containing 50 mM Tris-HCl (pH 7.4), 150 mM NaCl, 1 mM InsP_6_ (a commercially available surrogate for PP-InsPs^32^), 1% LMNG (Anatrace), 0.1% CHS (Anatrace) and 0.25% Soy Phospholipids (Sigma). The extraction were performed at 4 °C for 1.5 h, and the resulting solution was centrifuged at 23,000 g for 40 min. The supernatant was collected and incubated with anti-Flag G1 affinity resin (Genscript) at 4 °C for 40 min, further rinsed with 30 bed volumes of wash (W1) buffer containing 50 mM Tris-HCl (pH 7.4), 150 mM NaCl, 1 mM InsP_6_ and 0.02% GDN (Anatrace), and eluted by W1 buffer supplemented with 250 μg ml^-1^ Flag peptide (Genscript). The eluent was concentrated and further purified by size-exclusion chromatography (Superose-6 Increase 10/300 column, GE Healthcare) using a buffer containing 25 mM Tris-HCl (pH 8.0), 150 mM NaCl, 2 mM DTT, 1 mM InsP_6_ and 0.02% GDN. The peak fractions of XPR1 were collected and pooled to ∼4.5 mg ml^−1^ for cryo-EM grid preparation.

Sample used for the liposome-based transport assays were extracted from cultured cells by a buffer containing 50 mM Tris-HCl (pH 7.4), 150 mM NaCl and 1% n-Dodecyl-β-D-Maltopyranoside (DDM). Target proteins were further purified using anti-Flag G1 affinity resin and Superose-6 Increase 10/300 column in tandem, and prepared in the buffer containing 25 mM Hepes-Tris (pH 7.4), 150 mM NaCl, 2 mM DTT and 0.02% DDM for further proteoliposomes reconstitution.

### Cryo-EM grid preparation and data collection

3.5 μl aliquots of the purified protein was dispensed onto glow discharged holey carbon grid (Quantifoil Cu R1.2/1.3, 300 mesh). The gird was blotted with a Vitrobot Mark IV (ThemoFisher Scientific) using 3.5 s blotting time with 100% humidity at 8 °C, and was plunge-frozen in liquid ethane. The cryo-grid was transferred to 300 kV Titan Krios electron microscopes (Thermo Fisher) equipped with a GIF Quantum energy filter (slit width 20 eV) and a Gatan K3 Summit detector. EPU software (v2.9) was used for fully automated data collection. Micrographs were recorded in the super-resolution mode with a magnification of 81,000×. Each micrograph stack, which contains 32 frames, was exposed for 3.5 s with a total electron dose of 50 e^−^/Å^2^. MotionCor2 (v1.4.7)^53^ was used to perform beam-induced motion correction on cryo-EM images with binning factor of 2, resulting in a pixel size of 1.07 Å. The defocus value of each image was set to −1.2 to −2.2 μm and estimated by CTFFIND4 (v4.1.14)^54^.

### Cryo-EM data processing

A diagram of the procedures for data processing was described in Fig. S2. For the structure determination, 19,877 micrographs were manually selected from two datasets of 20,420 micrographs. A total of 24,071,915 particles were selected and extracted for 2D classification, out of which 23,126,090 particles were selected for 3D classification. After several rounds of 3D classification using the “multi-reference” approach, the 1,168,388 particles with the best class were re-extracted to their original size for 3D refinement, resulting in a cryo-EM density map with an overall resolution of 2.9 Å. This map clearly displays the transmembrane domain of XPR1, which forms dimers in the micelles. To preserve asymmetric features, C1 symmetry was employed during the 3D refinement. To search for and determine potential phosphate densities within XPR1 transmembrane domain, we further conducted structural heterogeneity analysis on these pooled particles using 3D Classification (BETA) in cyoSPARC and cryoDRGN^55,56^. These particles were mandatorily classified into 10 classes, with each particle belonging to only one class. The target resolution for each reconstruction was set at 3 Å, ensuring that each class was independent. Subsequently, a meticulous examination of the 3D-refined cryo-EM maps was conducted, resulting in the acquisition of a 2.9-Å cryo-EM map for XPR1 in the Pi-unbound form and three maps for the Pi-bound forms, with resolutions of 3.1, 3.1, and 3.3 Å, respectively. CryoSPARC (v4.5.1)^55^ and RELION (v5.0)^57^ were used for 2D classification, 3D classification and 3D refinement. Local resolution variations of the maps were estimated using Resmap (v1.1.4)^58^.

### Model building and refinement

The initial model of XPR1 was predicted from the Alphafold2^37^. We employed the ChimeraX software to dock this predicted model into the reconstructed cryo-EM map. The model was manually refined through iterative rounds of adjustments using COOT^59^. The residues of XPR1 transmembrane α-helices can be effectively constructed in the model. The phospholipid-like densities are assigned as phosphatidylcholine. Nonprotein densities were observed in one protomer of each dimer in the XPR1^Pi1^ and XPR1^Pi2^ cryo-EM maps, and in both protomers of the dimer in the XPR1^Pi3^ cryo-EM map. These densities were modeled with Pi ions. The obtained model was refined against the map using PHENIX^60^ in real space with secondary and geometry restrains. Model quality assessments were conducted via Molprobity scores^61^ and Ramachandran plots. Structural Figures were generated using ChimeraX (v1.6.1) and Pymol (v2.4.1).

### Transport assay

Liposomes (10 mg/mL) were prepared with E. coli total extract (Avanti Polar Lipids) in a reconstitution buffer containing 10 mM Hepes-Tris (pH 7.4) and 100 mM KCl. Preformed liposomes were dissolved with 1.3 % (w/v) DDM and mixed with purified XPR1 or variants in a protein-to-lipid ratio of 1:100 (w/w). Following incubation at 4 °C for 1.5 h, the DDM was removed by 3 additions of SM-2 bio-beads (Bio-Rad), incubated for 2h/2h/overnight, respectively. Prior to the start of the transport assay, the proteoliposomes were extruded using polycarbonate filter with a pore size of 200 nm (Whatman). 15 μl proteoliposomes containing 0.2-0.5 μg protein were diluted into 80 μl reconstitution buffer. Pi transport reactions were initiated by adding a KPi mixture at given concentration, which contains non-labeled KH_2_PO_4_ and [^32^P] KH_2_PO_4_ (3.7 MBq/ μmol; PerkinElmer) in a molar ratio of 29:1-3:1. The assays were performed at 37 °C, and terminated at given times by diluting tenfold with ice-cold stop buffer (10 mM Hepes-Tris, pH 7.4, 100 mM KCl and 5 mM non-labeled KH_2_PO_4_), followed by rapid filtration through nitrocellulose membrane (Millipore, 0.22 μm Triton-free MCE). The filters were subsequently washed with 2×5 ml ice-cold stop buffer, placed in 5 mL Optiphase HiSafe 3 scintillation fluid and counted after 14 h. Background was defined as the counts of parallel transport assays that were terminated at the beginning of the reaction. After subtracting the background, the amount of phosphate transported inside the proteoliposomes was quantified by comparing it to a standard curve of ^32^P KH_2_PO_4_. This standard curve was established by diluting sole ^32^P KH_2_PO_4_ in water and recording the corresponding radioactive counts. It thus provided a quantitative relationship between radioactive counts and the amount of ^32^P, which was used to quantify the amount of Pi taken up during our transport reactions. The protein contained in proteoliposomes were resolved by SDS-PAGE and quantified using ImageJ. The Pi transport activity is determined by measuring phosphate uptake into proteoliposomes containing proteins (pmol Pi/μg protein). The assays were independently performed a minimal of three times, each with technical triplicates, to generate an overall mean and s.d.

### Yeast Complementation

Corresponding genes were separately constructed into a PRS416-ADH vector and transformed into the yeast Pi transport-deficient mutant YP100 strain^41^. Transformants were selected on synthetic complete (SC-uracil) yeast medium (containing 2% galactose, 0.67% yeast nitrogen base without amino acids, 0.2% appropriate amino acids and 2% agar) at pH 5.4. The medium was incubated at 28 °C for 2-3 days, and transformants were verified by PCR. 2× YPD medium was used for transformants to cultivate. Mid-exponential phase cells were collected, washed two times with YNB (without phosphate) and resuspended to OD_600_=1 in water. Equal volumes of 5-fold serial dilutions were spotted on YNB medium (supplemented with 2% glucose, 7.5 mM KH_2_PO_4_, 2% agar and 7.5% amino acids mix, deficient uracil) at pH 5.4. Plates were incubated at 28 °C for 7 days to observe yeast growth.

### Molecular dynamic simulations

Atomistic models of XPR1 transmembrane domain were constructed using the structures of Pi-unbound XPR1 and Pi-bound XPR1^Pi1^. The missing loop (amino acids 432-445) was modeled using the AlphaFold2 prediction as a template. These models were then embedded in a POPC lipid bilayer and solvated in a cubic water box containing 0.15 M NaCl. The lipids identified in the cryo-EM density maps were also modeled as POPC. The dimensions of the box were 8.0 nm × 8.0 nm × 8.2 nm in the x, y, and z directions, respectively, totaling approximately 47,000 atoms. The transmembrane region was aligned in the lipid bilayer using the OPM (Orientations of Proteins in Membranes) webserver. Construction of the systems was facilitated by the CHARMM-GUI webserver^62^, followed by energy minimization using the steepest descent algorithm and a six-step equilibration process with gradual removal of position constraints. Production runs under semi-isothermal-isobaric (NPT) conditions were conducted using the CHARMM36m force field^63^ for proteins and lipids, along with the TIP3P model for water. The Pi ion in the XPR1^Pi^^1^ structure was modeled as H_2_PO_4_^-^. The protonation states of residues in the Pi binding sites were assigned according to their pKa values predicted by PROPKA3.1^64^ and further justified by examining the local environment. During the MD simulations, temperature was maintained at 310 K with a Nosé–Hoover thermostat and a coupling constant of 1 ps, and pressure was kept at 1.0 bar using the Parrinello–Rahman barostat and a time coupling constant of 5 ps. A switch function with a starting distance of 1.0 nm was employed for a van der Waals cut-off of 1.2 nm. Short-range electrostatic interactions were also truncated at 1.2 nm, while long-range electrostatic interactions were computed via the particle mesh Ewald decomposition algorithm with a 0.12-nm mesh spacing. For each system, two independent 1000 ns MD simulations were conducted using the GPU-accelerated version of Gromacs 2024.2^65^.

To accelerate Pi transport and enable the observation of transport events within feasible simulation timescales, a constant electrostatic field along the membrane normal was applied to simulate a transmembrane voltage of approximately 400 mV. Although this voltage is higher than typical physiological conditions (usually around -70 to -90 mV in cells), it is widely used in simulations of ion transporters. This value is considered high enough to drive the transport process effectively while remaining within a range that ensures the robustness and relevance of the simulation results. Trajectory analysis was performed using Gromacs gmx tools and PLUMED 2.8.2^66^. The water density map was analyzed with GROmaps^67^.

### Data availability

The EM maps and atomic models of Pi-unbound XPR1, Pi-bound XPR1^Pi1^, XPR1^Pi2^ and XPR1^Pi3^ have been deposited in the Electron Microscopy Data Bank and Protein Data Bank, with the accession numbers EMD-61138, EMD-61139, EMD-6140 and EMD-61141, and 9J4X.PDB, 9J51.PDB, 9J52.PDB and 9J53.PDB, respectively. Materials are available from the corresponding authors on request.

## Acknowledgments

We thank the Cryo-EM Center, the University of Science and Technology of China (USTC), for the EM facility support. We are grateful to Dr. Yongxiang Gao (USTC) for technical support during EM image acquisition. We thank the Center for Protein Research, and Dr. Jianbo Cao at the Public Laboratory of Electron Microscopy, Huazhong Agricultural University, for technical support. We thank the Information Technology Center and State Key Lab of CAD&CG, Zhejiang University, for computational support. We thank Prof. Michael Hothorn and Prof. Xuewen Cheng for helpful suggestions on the manuscript. This work was supported by the National Key Research and Development Program of China (2023YFF1000700), the National Natural Science Foundation of China (32071226 and 32371300), the Foundation of Hubei Hongshan Laboratory (2021HSZD011), the Fundamental Research Funds for the Central Universities (2662023PY001), the HZAU-AGIS Cooperation Fund (SZYJY2022022), and the Zhejiang Provincial National Science Foundation of China (No. LZ24C050003). Z.G. acknowledges the support of National Postdoctoral Program for Innovative Talents (BX2021108).

## Author contributions

Z.L. conceived and supervised the project. W.Z., Y.C. and Z.G. designed all experiments. W.Z. prepared samples. Y.C. performed transport assays. Z.G. determined the structures. Y..W. performed the MD simulations. M.T., Z.D., J.Z., M.C., J.Z. and Y.L. contributed to plasmids constructing and data collecting. Q.W., Y.L., D.Z., P.Y. and L.M. contributed in data analysis. Z.L. and Y.C. wrote the manuscript with help from all authors.

## Supplementary figures and table

**Fig. S1.**
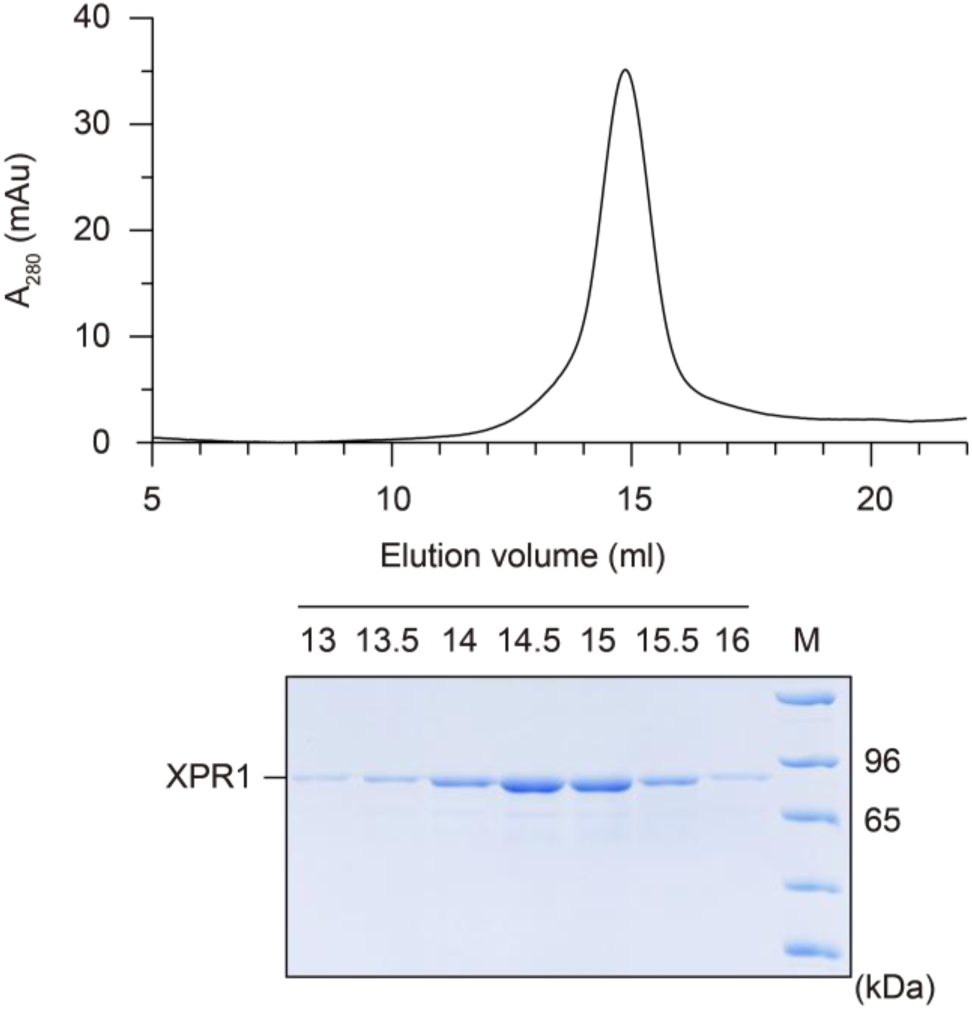
XPR1 purification for cryo-EM analysis. Representative size-exclusion chromatography (SEC) purification of XPR1 in the presence of InsP_6_. Fractions were resolved by 7% SDS-PAGE and the peak fractions were collected and concentrated to prepare cryo-EM grids.

**Fig. S2.**
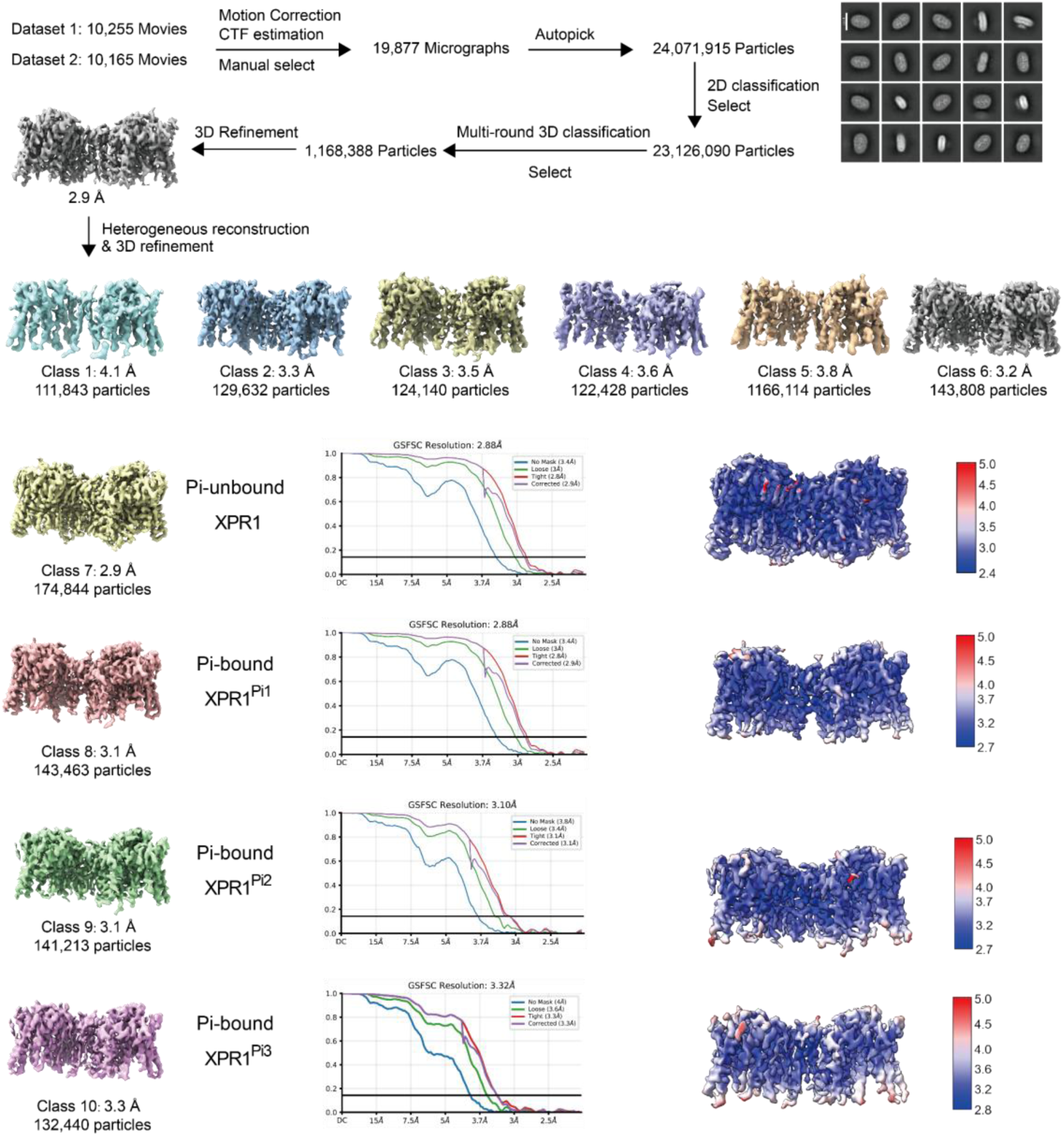
Cryo-EM data processing workflow of XPR1. The final reconstructed maps (left panel), gold standard Fourier shell correlation (FSC) curves (middle panel) and local resolution (right panel) are represented from the 3D refinement for the final reconstructions.

**Fig. S3.**
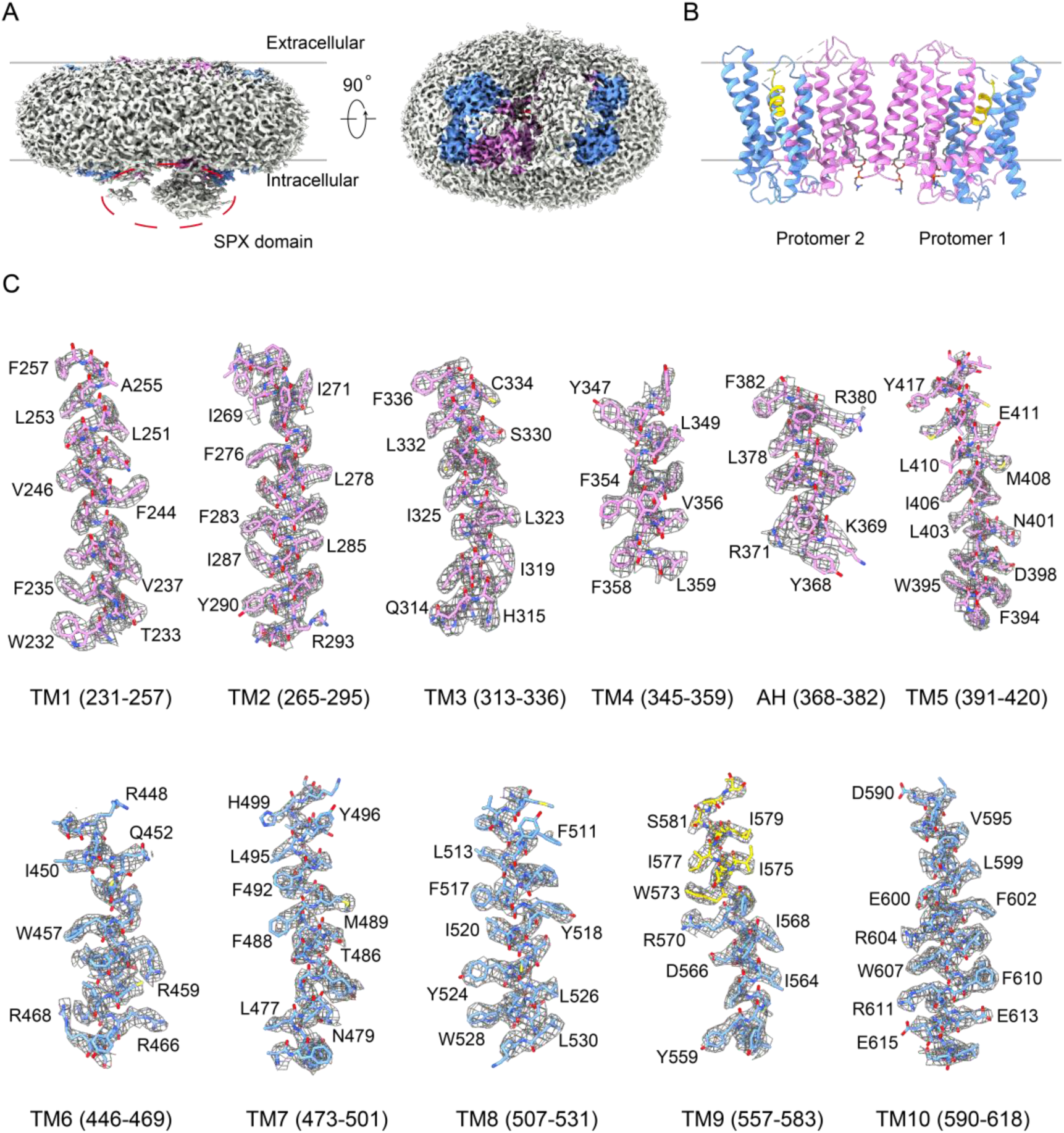
EM densities and representative elements of XPR1 (Pi-unbound). **A** EM density map with low display threshold reveals the signals of the cytoplasmic SPX domain and detergent. The transmembrane domain of XPR1 is colored in the same scheme as Fig. 1A in the main text. XPR1 presents as a dimer in the EM map. **B** Cartoon representation of the resolved dimeric transmembrane domain of XPR1. The two protomers adopt a similar structure with an overall RMSD of 0.26 Å . Protomer 1 (chain A) is used for further analysis and discussion in this manuscript. Lipid-like densities have been modeled using phosphatidylcholine molecules and are shown as sticks. **C** EM density maps for the indicated transmembrane α-helices. Map contour level = 1.0 in ChimeraX.

**Fig. S4.**
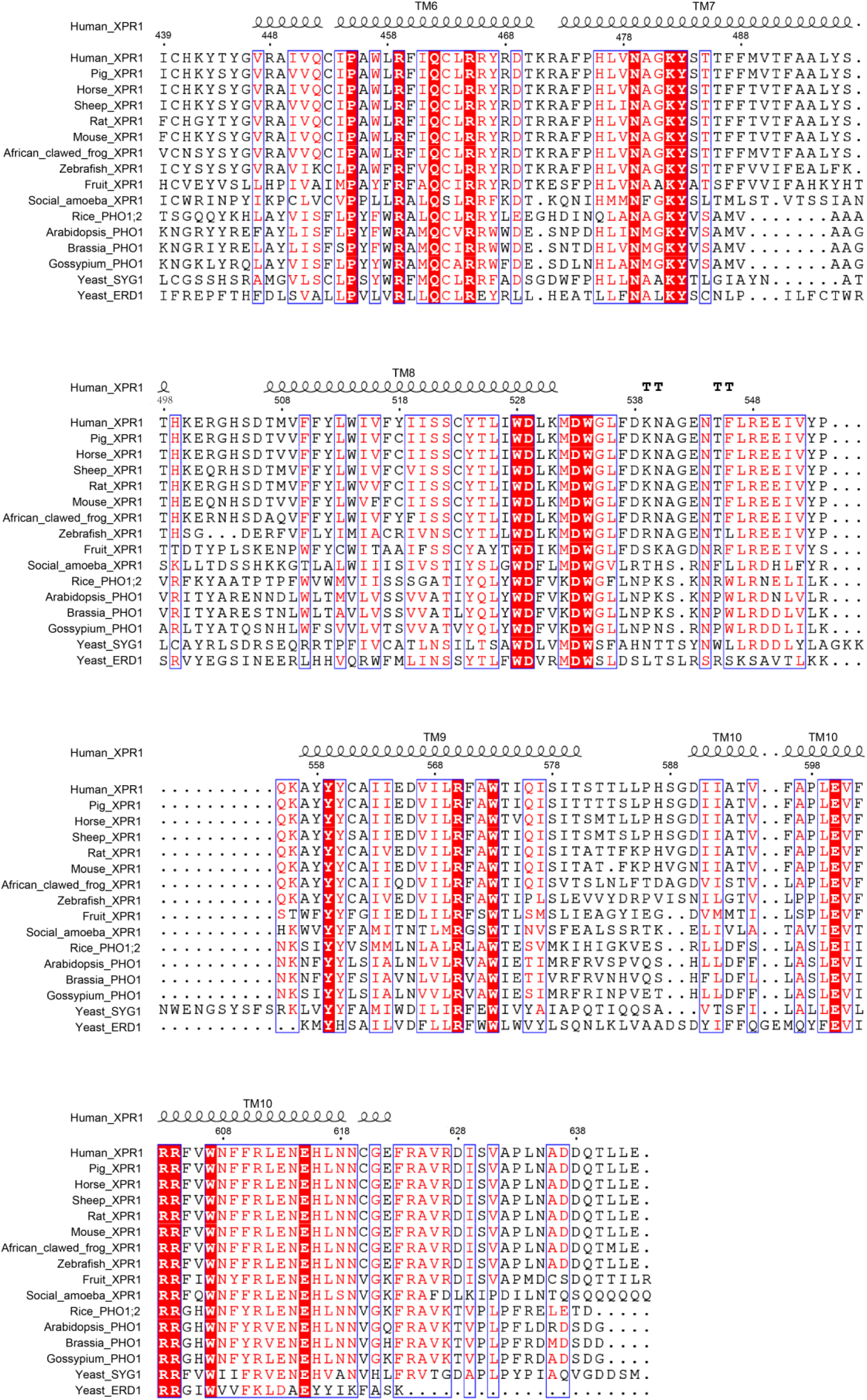
Sequence alignment of EXS domain-containing proteins. The sequence of EXS domain from human XPR1 is aligned with other orthologues. Structure-based alignment was performed by ESPript (3.0). The sequence identity is indicated by white letters against a red background, and the sequence of similarity over 90% is indicated by red letters. The secondary elements of EXS domain from human XPR1 are labeled at the top of the alignment.

**Fig. S5.**
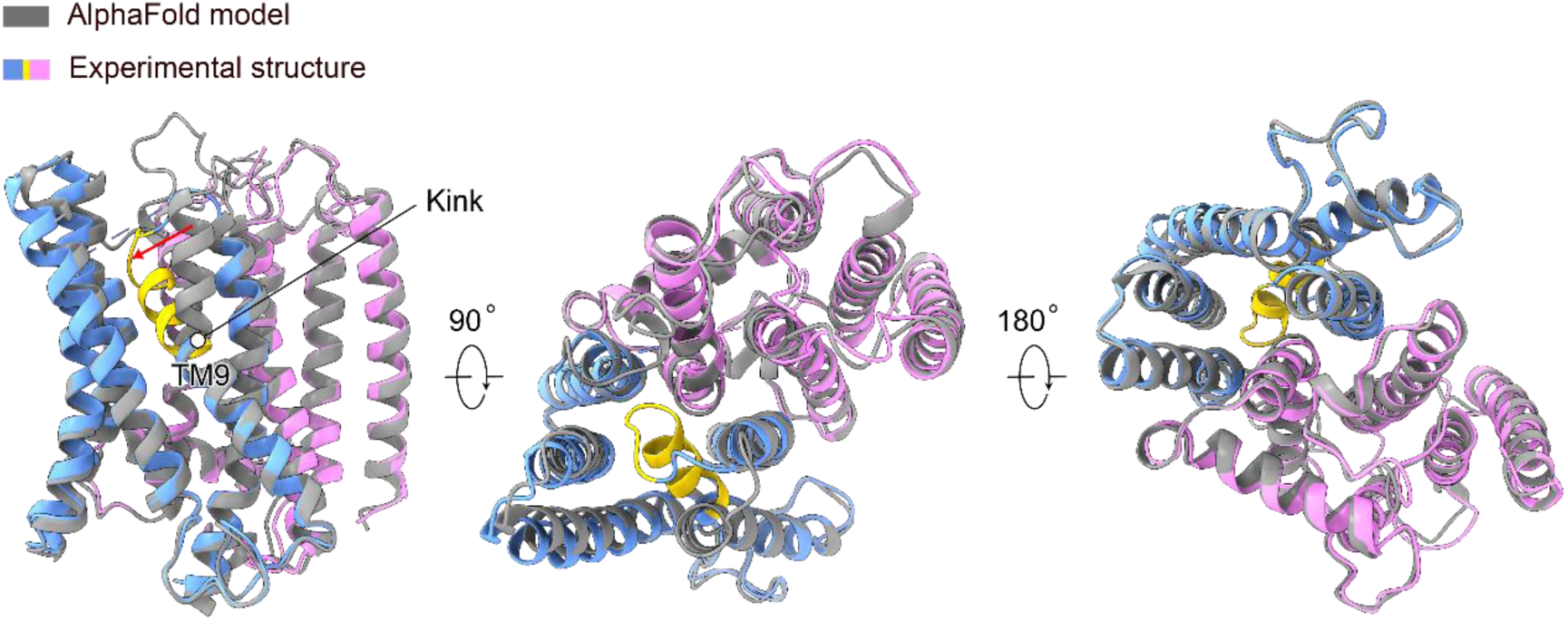
Comparison between the experimental structure of XPR1 transmembrane domain and the predicted model by AlphaFold2. The experimental structure is colored in the same scheme as Fig. 1A in the main text. The AlphaFold2 model (AF-Q9UBH6-F1) is superposed and colored in gray. A red arrow indicates the bending of the extracellular half of TM9 observed in the experimental structure.

**Fig. S6.**
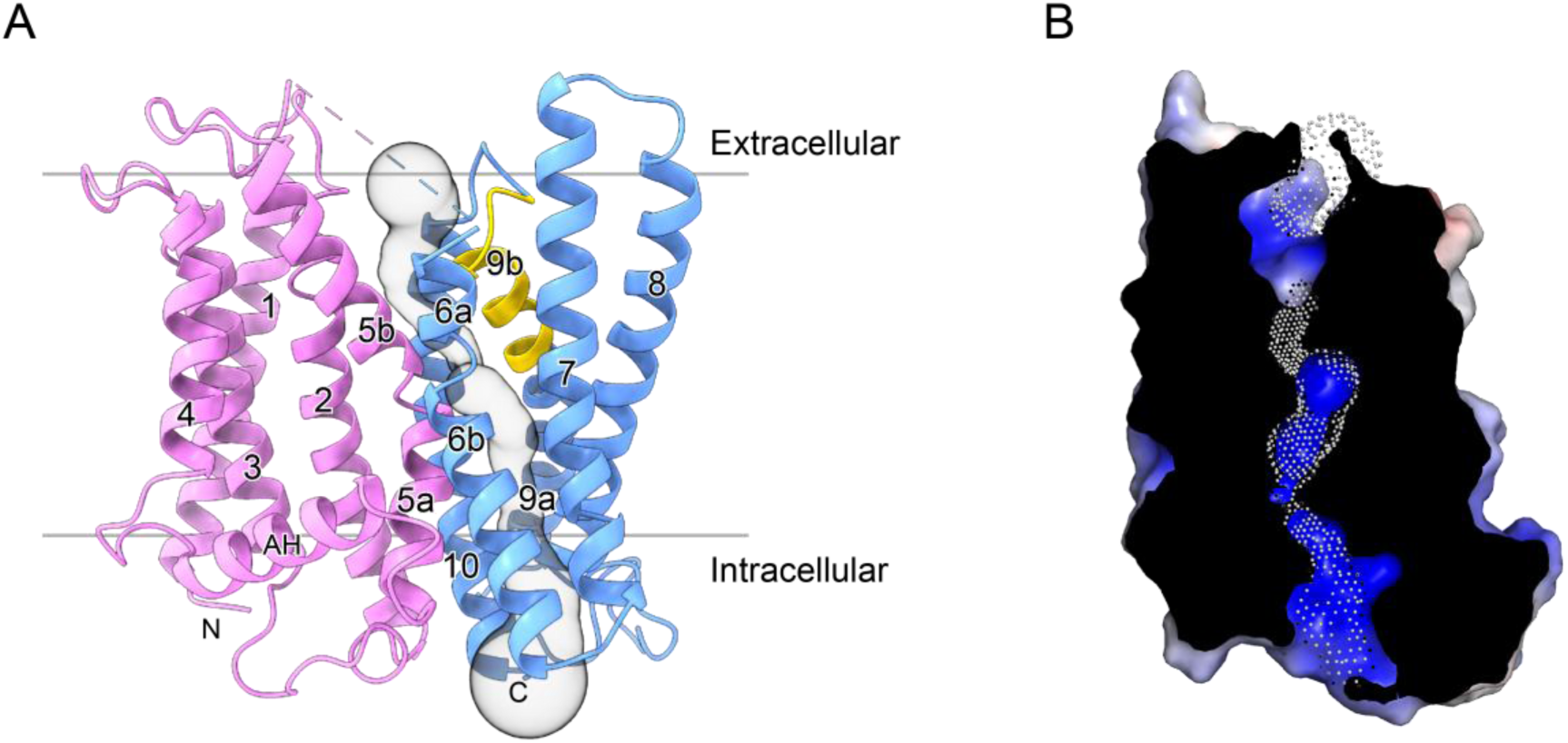
Transmembrane channel. **A** Solvent-accessible pathway (gray surface representation) within the XPR1 transmembrane domain. The structure is represented in the same scheme as Fig. 1A in the main text. **B** A cut-open electrostatic surface representation of the channel. It is colored in terms of electrostatic potential, and displayed in a scale from red (−5 kT/e) to blue (+5 kT/e). For clarity, only the surface of channel-forming transmembrane α-helices are represented. The dots indicate the solvent-accessible pathway.

**Fig. S7.**
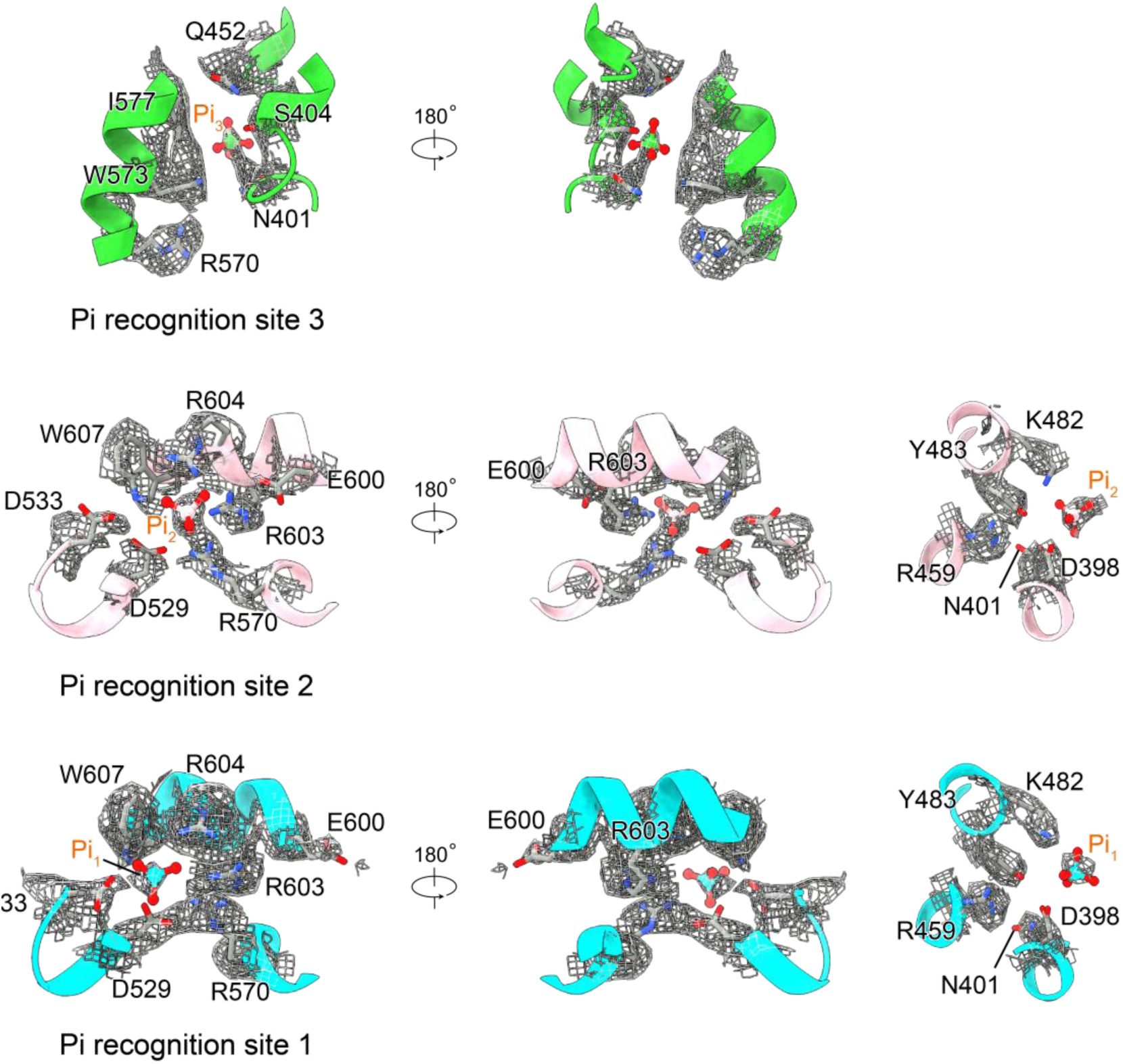
EM density maps of the Pi recognition sites 1-3. This figure corresponds to Fig. 2D-F in the main text. The densities derived from XPR1^Pi1^, XPR1^Pi2^ and XP1^Pi3^ structures are represented with contour levels of 0.841, 0.655 and 0.586, respectively, in ChimeraX.

**Fig. S8.**
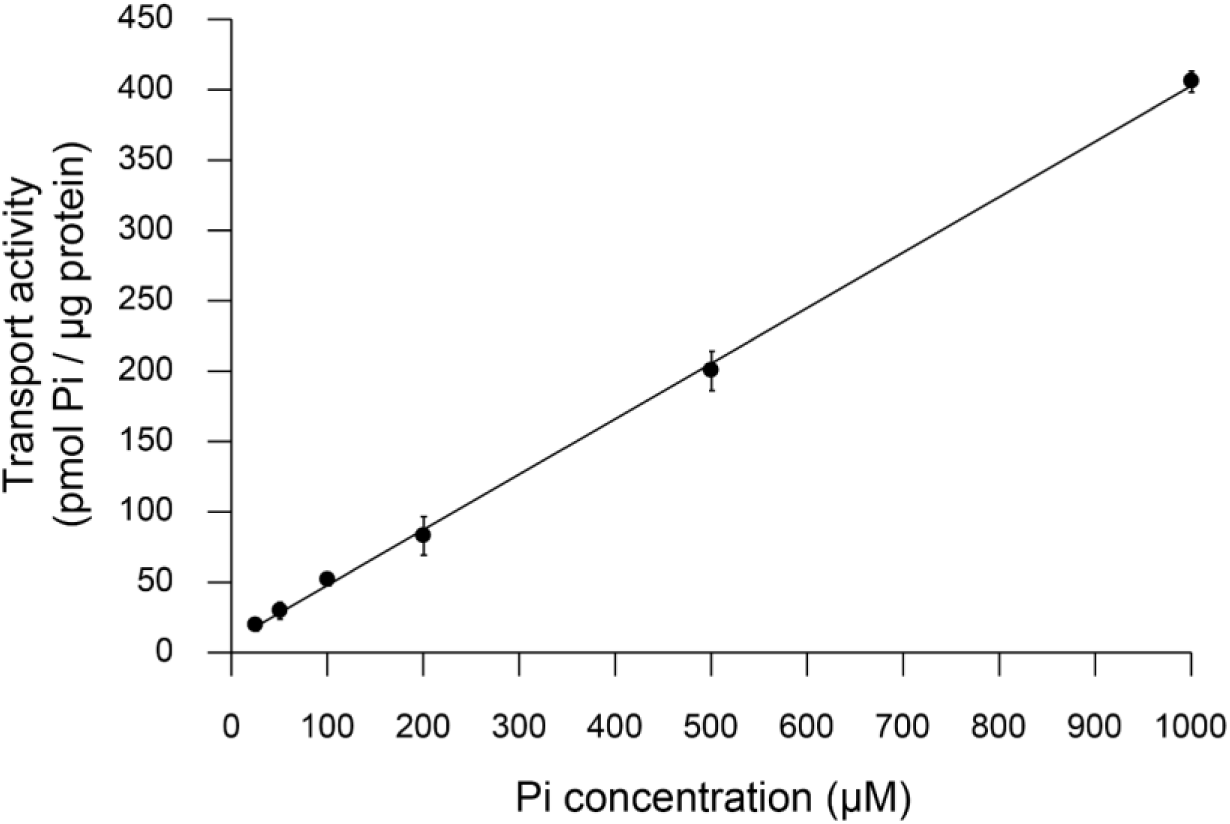
Proteoliposome-based Pi transport assay of XPR1. The transport reactions were allowed for 6 minutes with external Pi concentrations of 25, 50, 100, 200, 500 and 1000 μM, respectively. The data presented are the average of three independent assays, each with technical triplicates. The error bars indicates the SD.

**Fig. S9.**
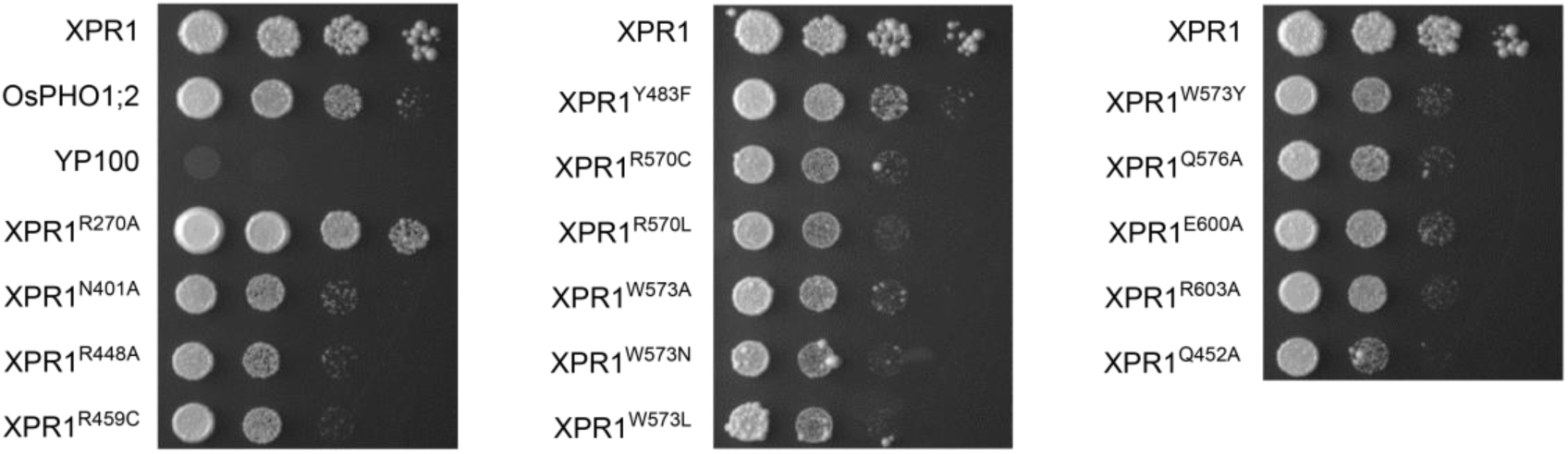
Complementation assays of the yeast mutant YP100. All yeast cells were grown on YNB medium for 7 days, and equal volumes of 5-fold serial dilutions were spotted onto the medium. The experiments were repeated independently at least three times with similar results.

**Fig. S10.**
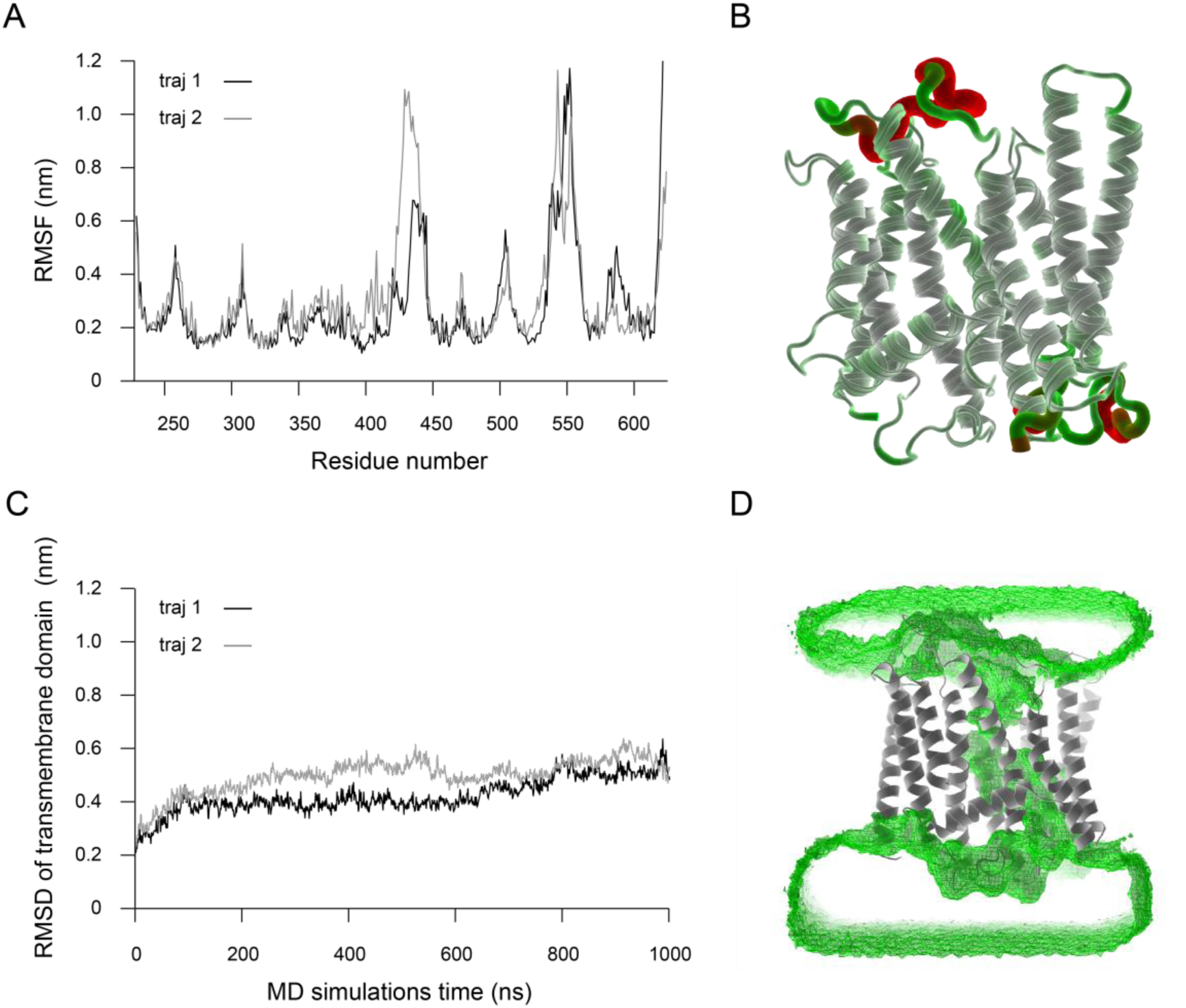
MD simulations of the cryo-EM structure of Pi-unbound XPR1. **A** RMS fluctuations of XPR1 residues during the simulations, calculated from two independent 1000-ns MD trajectories. **B** RMSF values per residue mapped onto the XPR1 structure from the final frame of the MD simulations. **C** RMSD trajectories of the heavy atoms in the transmembrane helices of XPR1 relative to the energy-minimized cryo-EM structure of Pi-unbound XPR1. **D** Consistent hydration observed in the transmembrane pore of XPR1 during one of the MD simulations, with average water density shown as green meshes.

**Fig. S11.**
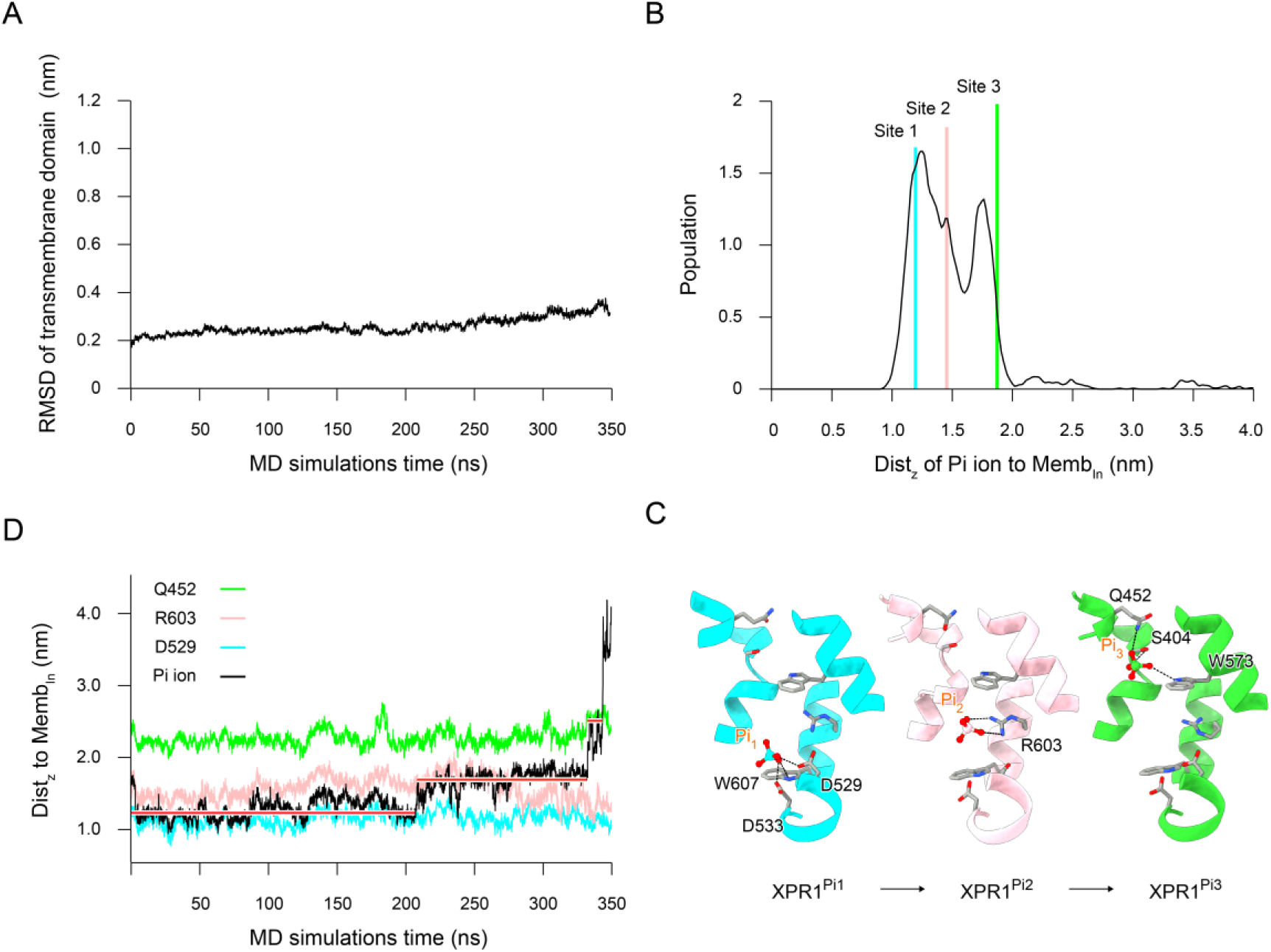
MD simulations and analysis of the cryo-EM structure of Pi-bound XPR1^Pi1^. **A** RMSD trajectories of the heavy atoms in the transmembrane helices of XPR1 relative to the energy-minimized cryo-EM structure of Pi-bound XPR1^Pi1^. **B** Distribution of the distance (Dist_z_) between the Pi ion and the center of mass of the phosphate head groups of lipids in the inner leaflet along the membrane’s normal direction (Z-axis). The positions of the three Pi recognition sites observed in the cryo-EM structures of XPR1^Pi1^, XPR1^Pi2^ and XPR1^Pi3^ are marked with cyan, pink, and green lines, respectively. **C** Side view of the channel highlighting specific residues involved in Pi coordination at the recognition sites 1, 2, and 3. This figure relates to Fig. 2D-F in the main text and focuses on site-specific interactions for clarity. **D** Correlation between the positions of the Pi ion and representative site-specific residues during the simulations. The red lines indicate the stepwise interactions with D529, R603, and Q452.

**Table S1.**
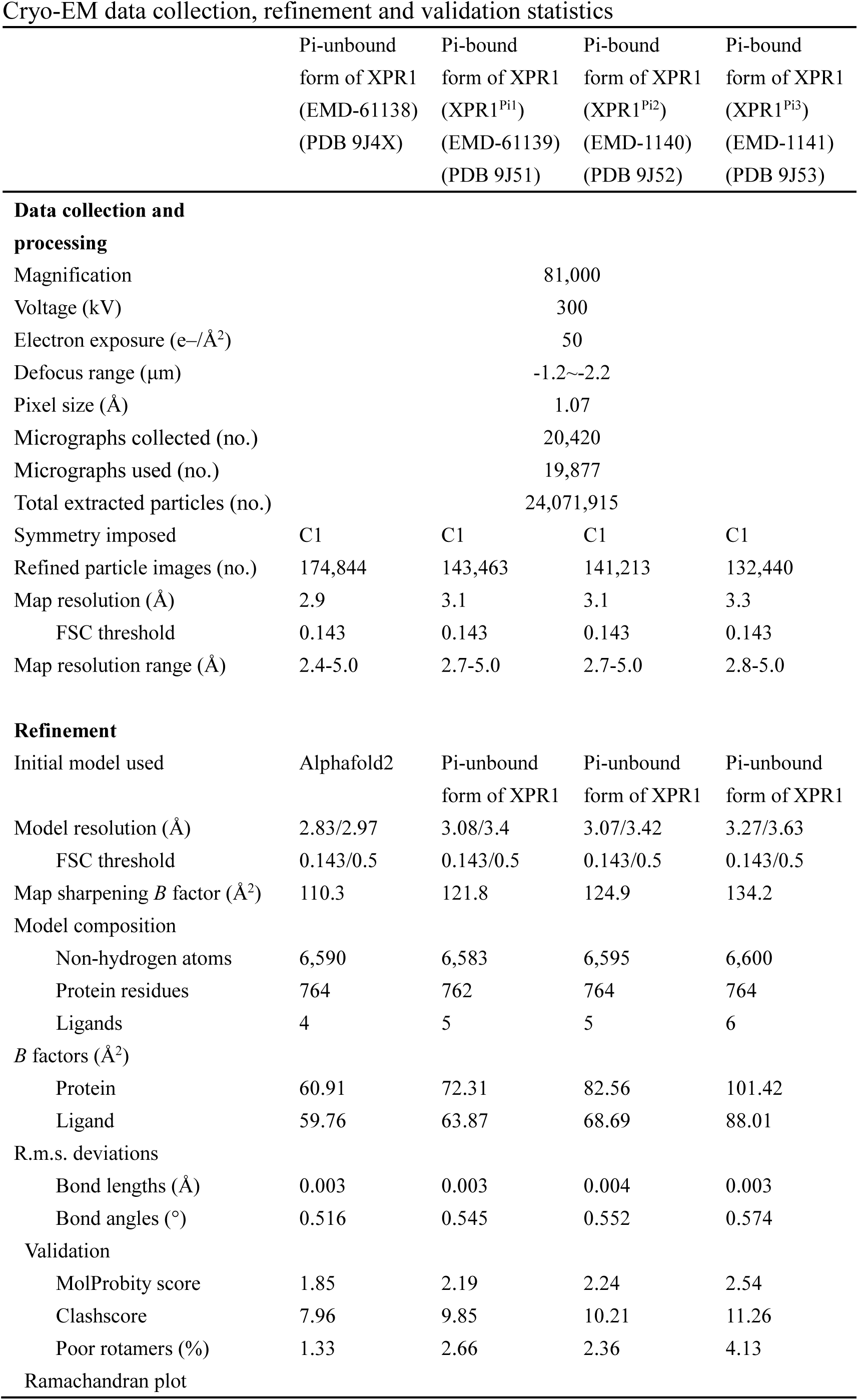
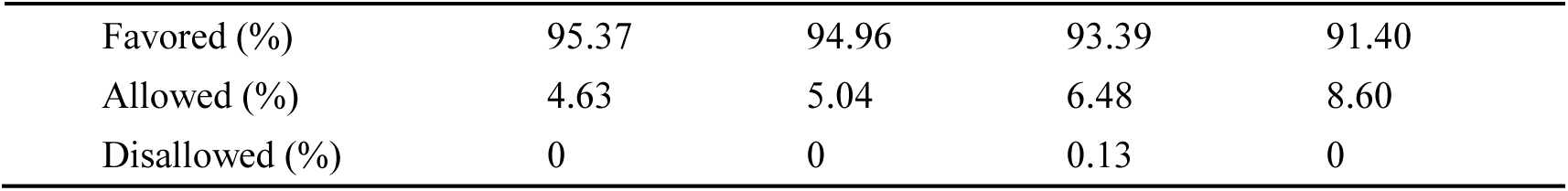
Cryo-EM data collection, refinement and validation statistics.

|  | Pi-unbound<br>form of XPR1<br>(EMD-61138)<br>(PDB 9J4X) | Pi-bound<br>form of XPR1<br>(XPR1<sup>Pi1</sup>)<br>(EMD-61139)<br>(PDB 9J51) | Pi-bound<br>form of XPR1<br>(XPR1<sup>Pi2</sup>)<br>(EMD-1140)<br>(PDB 9J52) | Pi-bound<br>form of XPR1<br>(XPR1<sup>Pi3</sup>)<br>(EMD-1141)<br>(PDB 9J53) |
| --- | --- | --- | --- | --- |
| **Data collection and processing** |  |  |  |  |
| Magnification |  |  | 81,000 |  |
| Voltage (kV) |  |  | 300 |  |
| Electron exposure (e-/Å<sup>2</sup>) |  |  | 50 |  |
| Defocus range (μm) |  |  | -1.2~-2.2 |  |
| Pixel size (Å) |  |  | 1.07 |  |
| Micrographs collected (no.) |  |  | 20,420 |  |
| Micrographs used (no.) |  |  | 19,877 |  |
| Total extracted particles (no.) |  |  | 24,071,915 |  |
| Symmetry imposed | C1 | C1 | C1 | C1 |
| Refined particle images (no.) | 174,844 | 143,463 | 141,213 | 132,440 |
| Map resolution (Å) | 2.9 | 3.1 | 3.1 | 3.3 |
| FSC threshold | 0.143 | 0.143 | 0.143 | 0.143 |
| Map resolution range (Å) | 2.4-5.0 | 2.7-5.0 | 2.7-5.0 | 2.8-5.0 |
| **Refinement** |  |  |  |  |
| Initial model used | Alphafold2 | Pi-unbound<br>form of XPR1 | Pi-unbound<br>form of XPR1 | Pi-unbound<br>form of XPR1 |
| Model resolution (Å) | 2.83/2.97 | 3.08/3.4 | 3.07/3.42 | 3.27/3.63 |
| FSC threshold | 0.143/0.5 | 0.143/0.5 | 0.143/0.5 | 0.143/0.5 |
| Map sharpening *B* factor (Å<sup>2</sup>) | 110.3 | 121.8 | 124.9 | 134.2 |
| Model composition |  |  |  |  |
| Non-hydrogen atoms | 6,590 | 6,583 | 6,595 | 6,600 |
| Protein residues | 764 | 762 | 764 | 764 |
| Ligands | 4 | 5 | 5 | 6 |
| *B* factors (Å<sup>2</sup>) |  |  |  |  |
| Protein | 60.91 | 72.31 | 82.56 | 101.42 |
| Ligand | 59.76 | 63.87 | 68.69 | 88.01 |
| R.m.s. deviations |  |  |  |  |
| Bond lengths (Å) | 0.003 | 0.003 | 0.004 | 0.003 |
| Bond angles (°) | 0.516 | 0.545 | 0.552 | 0.574 |
| Validation |  |  |  |  |
| MolProbity score | 1.85 | 2.19 | 2.24 | 2.54 |
| Clashscore | 7.96 | 9.85 | 10.21 | 11.26 |
| Poor rotamers (%) | 1.33 | 2.66 | 2.36 | 4.13 |
| Ramachandran plot |  |  |  |  |
| Favored (%) | 95.37 | 94.96 | 93.39 | 91.40 |
| Allowed (%) | 4.63 | 5.04 | 6.48 | 8.60 |
| Disallowed (%) | 0 | 0 | 0.13 | 0 |

**Table S2.**
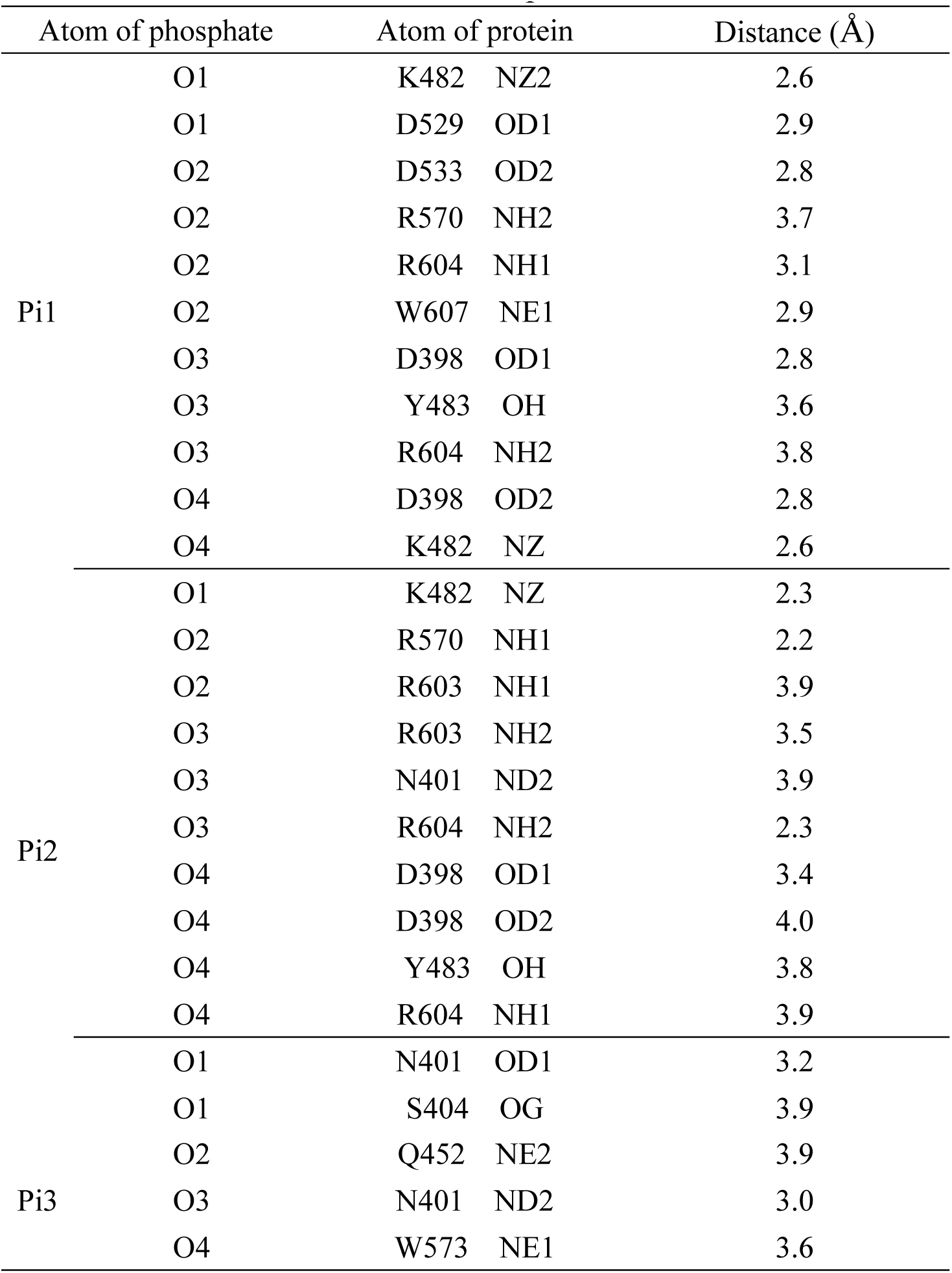
Plausible interactions between the Pi ions and protein.

